# Self-regulating living material with temperature-dependent light absorption

**DOI:** 10.1101/2023.03.11.532239

**Authors:** Lealia L. Xiong, Michael A. Garrett, Julia A. Kornfield, Mikhail G. Shapiro

## Abstract

Engineered living materials (ELMs) exhibit desirable characteristics of the living component, including growth and repair, and responsiveness to external stimuli. *Escherichia coli* are a promising constituent of ELMs because they are very tractable to genetic engineering, produce heterologous proteins readily, and grow exponentially. However, seasonal variation in ambient temperature presents a challenge in deploying ELMs outside of a laboratory environment, because *E. coli* growth rate is impaired both below and above 37°C. Here, we develop a genetically-encoded mechanism for autonomous temperature homeostasis in ELMs containing *E. coli* by engineering circuits that control the expression of a light-absorptive chromophore in response to changes in temperature. We demonstrate that below 36°C, our engineered *E. coli* increase in pigmentation, causing an increase in sample temperature and growth rate above non-pigmented counterparts in a model planar ELM. On the other hand, above 36°C, they decrease in pigmentation, protecting their growth compared to bacteria with temperature-independent high pigmentation. Integrating our temperature homeostasis circuit into an ELM has the potential to improve living material performance by optimizing growth and protein production in the face of seasonal temperature changes.

## INTRODUCTION

The field of engineered living materials (ELMs) aims to confer desirable properties of living cells and naturally-occurring biomaterials to designed materials either by incorporating biological cells into a synthetic material, or through the formation of a material by living cells and their synthesized biopolymers^[1–3]^. These properties include self-assembly, self-healing, and sensing and responding to signals. The cells of microbial species including yeast^[4]^, cyanobacteria^[5]^ and *Bacillus subtilis*^[6,7]^ have been used in ELMs, with *Escherichia coli* an especially common focus of ELM researchers^[2,8–10]^ because of its extensive characterization and tractability for genetic engineering^[11]^.

While *E. coli* is a laboratory workhorse, it normally lives in the intestines of warm-blooded animals, where the host maintains a stable temperature of about 37°C^[12,13]^. However, outside of the laboratory or intestinal environment, cells are exposed to fluctuations in temperature, nutrients, and moisture, all of which affect their viability and growth rate. Indeed, *E. coli* is known to live in freshwater and soil only in tropical ecosystems, which maintain an appropriate temperature, nutrient level, and humidity^[12]^. In the laboratory, the minimum temperature for growth of *E. coli* is about 7.5°C in minimal media^[14]^, but even a decrease in temperature from 37°C to 25°C reduces the rate of growth of *E. coli* cultures in exponential phase by about 38%^[15]^. The reduction in growth rate with temperature correlates with reduced ability to synthesize protein. On the other hand, exposure to temperature above 37°C is also deterimental to *E. coli*, with growth inhibited above about 44°C, due to protein instability at high temperatures^[16]^. Thus, exposure to nonideal temperature limits the potential for environmental or building materials applications for an ELM in which *E. coli* is a living component.

Here, we develop a genetic circuit that senses temperature and turns on the formation of a light-absorptive pigment below 36°C. We demonstrate that this circuit enables *E. coli* growing in dense patches under simulated sunlight to improve its growth rate over unpigmented *E. coli* at a suboptimal environmental temperature of 32°C by capturing light and warming up. At the same time, when growing at 42°C, an above-optimal temperature, our engineered *E. coli* remains unpigmented and grows faster than bacteria with temperature-independent pigmentation.

## RESULTS

*E. coli* within ELMs in outdoor applications, such as building exteriors, will be exposed to changes in ambient temperature, which will challenge their ability to grow and produce proteins of interest. For *E. coli*, the optimal growth temperature is 37°C. We propose that when the ambient temperature drops below the optimum, the cells should produce a light-absorptive pigment, allowing them to warm up when exposed to sunlight, whereas at or above the optimum, they should remain unpigmented, to avoid overheating **(Figure 1a)**.

**Figure 1.**
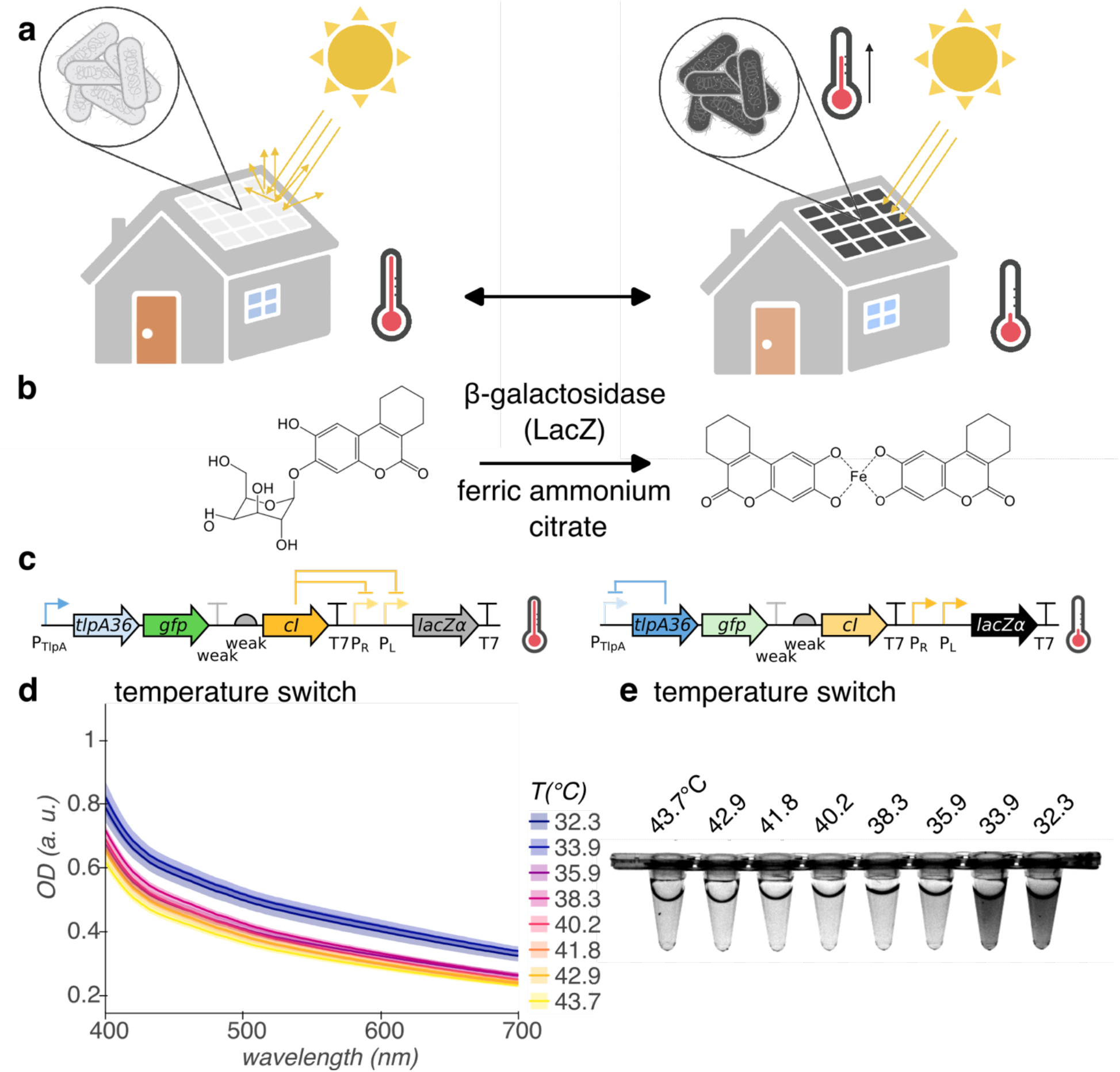
Cold-activated production of light-absorptive pigment for *E. coli-containing* ELMs. **(a)** Illustration of ELM used as building material. At ambient temperature greater than or equal to optimum for growth, *E. coli* remain colorless (left). However, at ambient temperature less than optimal, *E. coli* express light-absorptive pigment, warming under illumination by the sun to recover growth rate (right). Illustration created with BioRender.com. **(b)** Genetically-encodable light-absorptive pigment system. β-galactosidase cleaves S-gal at the glycosidic bond, exposing the esculetin group, which coordinates with ferric iron to form a black pigment. **(c)** Circuit diagram of temperature switch construct for low-temperature pigmentation, with state of regulation arcs indicated at high and low temperature. Genes, left to right: *tlpA36, gfp, cI, LacZα*. **(d, e)** Visible light absorption spectra (**d**) and representative white light transillumination image (**e**) of cultures of *E. coli* containing the temperature switch construct after 24 h growth in pigment-induction media at temperatures ranging from 43.7°C to 32.3°C. *n* = 4 biological replicates; shading represents +/− standard error of the mean.

We repurposed a chemical compound designed to enable the use of ß-galactosidase as a gene reporter, 3,4-cyclohexenoesculetin-β-D-galactopyranoside (S-gal), as our genetically-encodable pigment **(Figure 1b)**. S-gal, which is yellow in solution, consists of galactose linked to 3,4-cyclohexenoesculetin by a glycosidic bond. β-galactosidase hydrolyzes this bond, which frees the esculetin group to complex with ferric iron (provided in the growth medium as ferric ammonium citrate, which is brown in solution) to form a black light-absorptive pigment^[17]^.

We used the TlpA36 transcriptional repressor as a genetically-encodable sensor with a temperature threshold of 36°C^[18]^ and constructed a gene circuit to invert its action, enabling *E. coli* to turn on enzymatic pigment production below 36°C **(Figure 1c)**. At these temperatures, TlpA36 represses expression of CI repressor from the **P**_TlpA_ promoter. The LacZα peptide of β-galactosidase, which presents a much lower metabolic burden to the cell than the complete enzyme, is expressed from the CI-regulated **P**_R_-**P**_L_ tandem promoter. Above 36°C, TlpA36 loses repressor function, so CI repressor is expressed from the **P**_TlpA_ promoter and represses expression of LacZα. The concentration of CI repressor is tuned down by placing a weak terminator and weak ribosome binding site upstream. The complementary LacZω peptide is induced from the genome of *E. coli* DH10B by addition of isopropyl β-D-1-thiogalactopyranoside (IPTG) to the growth medium. mWasabi (GFP) serves as a marker of high temperature.

We measured the visible light absorbance of planktonic *E. coli* DH10B containing our temperature switch construct after growth in pigment induction media (containing S-gal, ferric ammonium citrate, and IPTG) at temperatures ranging from 43.7°C to 32.3°C **(Figure 1d,e)**. Cultures of *E. coli* containing the temperature switch construct increase in absorbance across the visible light spectrum from 400 – 700 nm below 35.9°C, compared to at and above 35.9°C.

After constructing the temperature switch for expression of pigment below 36°C and demonstrating its function in liquid culture, we grew *E. coli* containing the construct in a dense, centimeter-scale patch format to simulate the environment of an ELM. We suction-coated a suspension of cells grown to saturation overnight at 38°C onto a polycarbonate membrane backing and placed it on a media-saturated melamine foam growth substrate **(Figure 2a)**. We placed the samples in a home-built illuminated growth chamber and monitored the relative temperature of the samples compared to the melamine foam using a lightweight 32×24 thermal IR sensor array **(Figure 2b)**. We placed the sensor array on a motorized arm that retracts when not imaging to avoid casting a shadow. After 48 h growth in illuminated conditions at 42°C, patches of *E. coli* containing our temperature switch construct produce little pigment, whereas at 32°C, they become nearly opaque when imaged with white light transillumination **(Figure 2c, Supplementary Figure S4)**.

**Figure 2.**
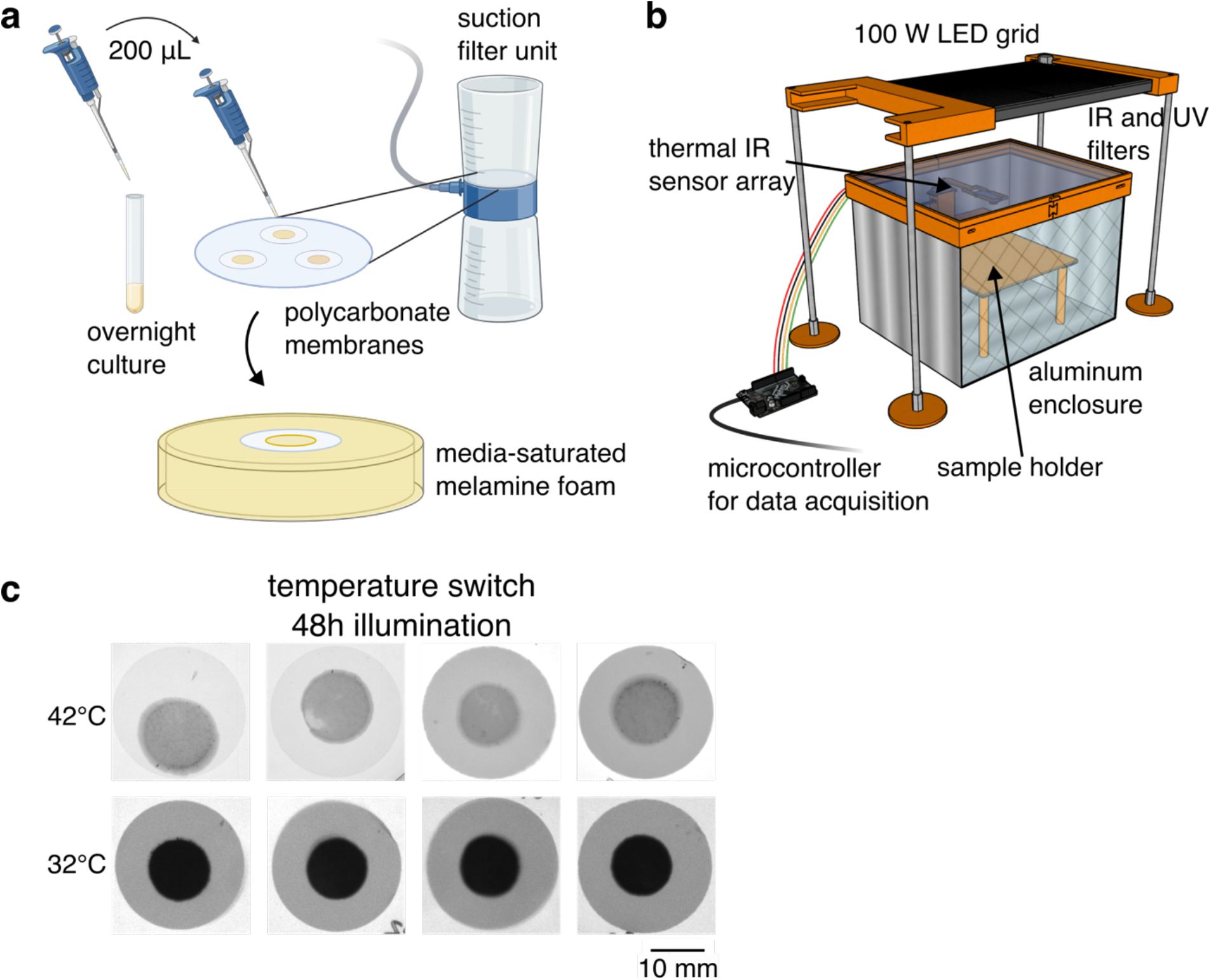
Temperature-dependent pigmentation in a model ELM. **(a)** Schematic of formation of dense, centimeter-scale patches of *E. coli* to simulate the ELM environment. We grew *E. coli* overnight to saturation in liquid medium. For each patch, we transferred 200 uL of culture to track-etched polycarbonate membranes (25 mm diam., 0.2 um pores) and applied suction to coat the cells onto the membranes, forming a dense patch. We then transferred the coated membranes to a melamine foam substrate saturated with liquid media for growth. **(b)** Schematic of illuminated growth chamber. We used a 100W white light LED to expose *E. coli* patches to illumination and monitored the temperature using a 32×24 array of thermal IR sensors. The sensor array is attached to a motorized arm and retracts when not imaging to avoid shadowing the samples. **(c)** Patches of *E. coli* containing the temperature switch construct on polycarbonate membranes after 48 h growth in the illuminated growth chamber with pigment-induction media at 42°C and 32°C. Parts of figure created with BioRender.com.

We tested the ability of turning on pigmentation to confer protection against slowed growth at 32°C with and without illumination by comparing the growth rate of *E. coli* containing our temperature switch construct against the growth rate of *E. coli* containing a control construct encoding mWasabi under the control of TlpA36^[18]^ (unpigmented control). While patches containing our temperature switch construct express pigment at this temperature, the control patches have no mechanism for pigment production and remain translucent **(Figure 3a)**. Due to the low spatial resolution of the sensor array, the precise temperature of the samples cannot be determined; however, the pigmented patches clearly warm above the background temperature **(Figure 3b)**.

**Figure 3.**
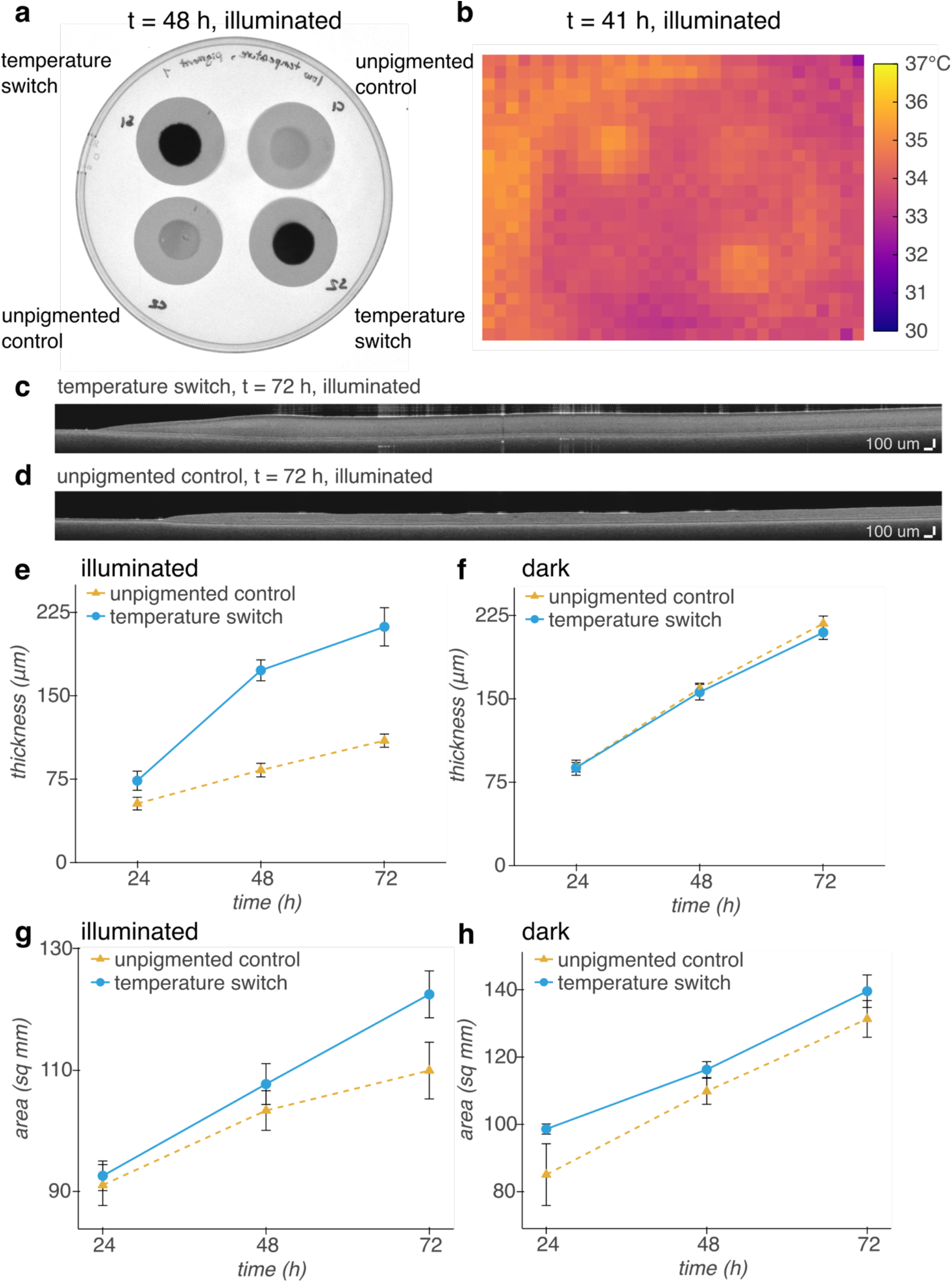
Cold-induced pigment improves the growth of dense patches of *E. coli* under illumination at 32°C. **(a)** White light transillumination image of patches of *E. coli* containing either our temperature switch construct or an unpigmented control construct encoding heat-inducible GFP after transferring to agar for imaging at 48 h. **(b)** Thermal IR image of patches inside illuminated growth chamber at 41 h. **(c, d)** Representative OCT images of patches of *E. coli* containing our temperature switch construct **(c)** or the unpigmented control construct **(d)**. **(e, f)** Thickness of patches grown under illumination **(e)** or in a dark incubator **(f)** over time. The slower rate of evaporation without illumination allows for thicker growth overall than with illumination. **(g, h)** Area of patches grown under illumination **(g)** or in a dark incubator **(h)** over time. *n* — 4 biological replicates; error bars represent +/− standard error of the mean.

We monitored the growth of the patches via their thickness, quantified from optical coherence tomography cross-sectional images, and their area, quantified from white light transillumination images. With illumination, the patches containing the temperature switch construct grow to twice the thickness of patches containing the control construct over the course of 72 h **(Figure 3c-e)**, whereas without illumination, the patches grow to the same thickness at each timepoint **(Figure 3f)**. In addition, with illumination, the temperature switch allows the patches to grow in area linearly with time, while the rate of increase in area of unpigmented patches slows over time **(Figure 3g)**. On the other hand, without illumination, patches containing both constructs grow in area linearly at approximately the same rate **(Figure 3h)**. These results suggest that expressing pigment at 32°C using the temperature switch construct gives *E. coli* in a dense patch format a growth advantage under illumination compared with unpigmented patches.

While pigmentation is advantageous at below-optimal ambient temperature, we hypothesized that it would be deleterious at above-optimal temperature by exacerbating overheating. To test this hypothesis, we compared the growth at 42°C of patches of *E. coli* containing our temperature switch construct against the growth rate of *E. coli* containing a control construct encoding IPTG-inducible LacZα (pigmented control). After 48 h, the patches containing our temperature switch construct lack pigmentation and appear translucent, while the control patches turn black and opaque, warming above the background temperature **(Figure 4a,b)**.

**Figure 4.**
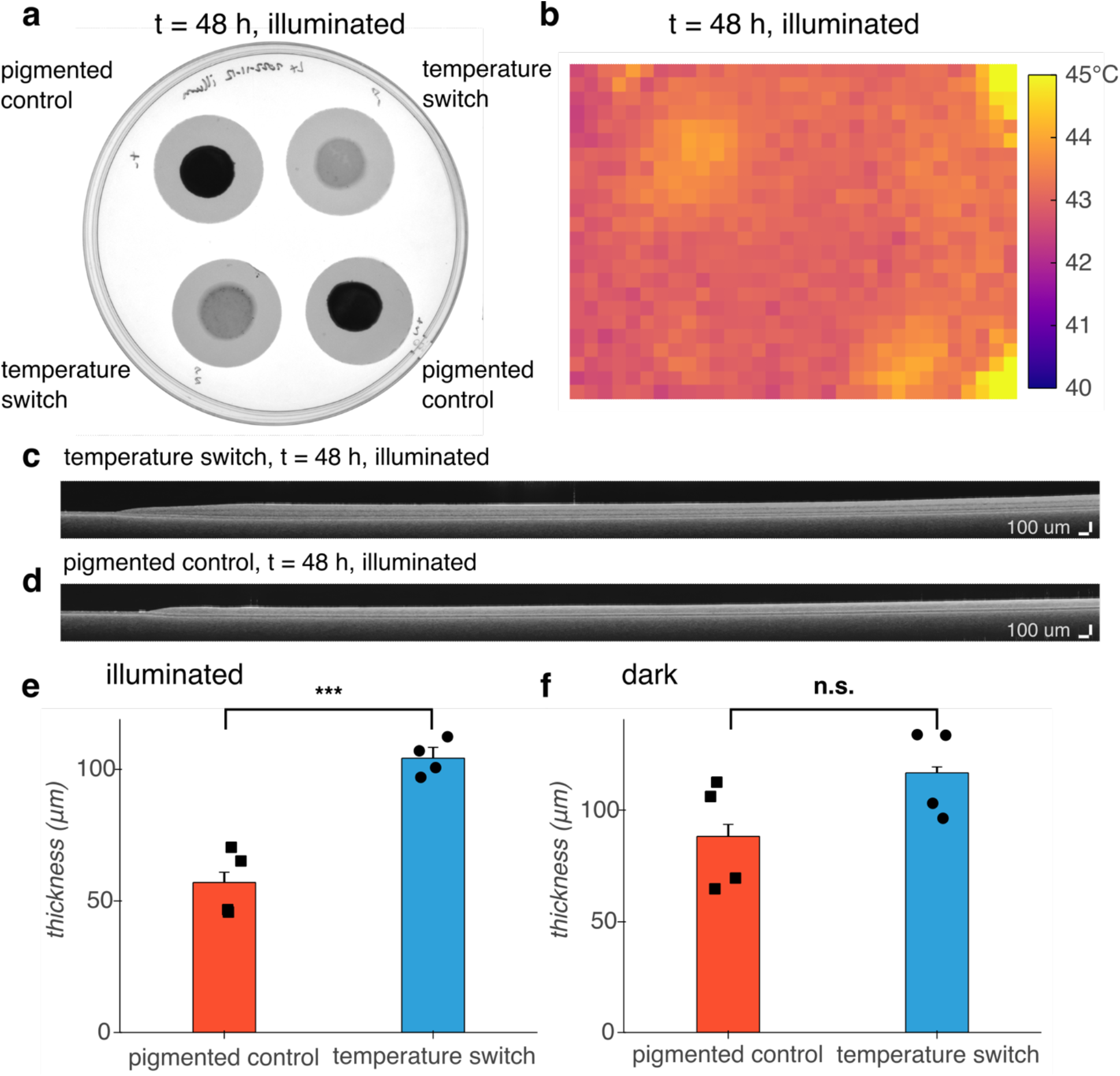
Turning off pigmentation above 36°C improves the growth of dense patches of *E. coli* under illumination at 42°C compared to a pigmented control. **(a)** White light transillumination image of patches of *E. coli* containing either the temperature switch construct or a pigmented control construct encoding IPTG-inducible LacZα after transferring to agar for imaging at 48 h. **(b)** Thermal IR image of patches inside illuminated growth chamber at 48 h. **(c, d)** Representative OCT images of patches of *E. coli* containing our temperature switch construct **(c)** or the pigmented control construct **(d)**. **(e, f)** Thickness of patches grown under illumination (p = 0.0006) **(e)** or in a dark incubator (p = 0.1214) **(f** at 48 h. *n* = 4 biological replicates; error bars represent +/− standard error of the mean. P-values calculated using a two-tailed unpaired *t*-test.

To avoid exposing the samples to laboratory room temperatures of 16 - 25°C by handling them during the experiment, we measured the area and thickness of the patches only at the endpoint of 48 h. The patches containing the temperature switch were thicker than the patches containing control construct under illumination **(Figure 4c-e)**. Without illumination, the patch thickness was not statistically different (p = 0.1214), although there was a trend toward greater thickness for the switch, potentially due to the burden of protein overexpression from the IPTG-induced control construct. Thus, turning off expression of pigment at 42°C using the temperature switch construct confers an advantage on *E. coli* in a dense patch format grown under illumination compared with temperature-independent, chemically-induced pigmentation.

## DISCUSSION

This work establishes a proof of concept for a genetically-encoded mechanism for supporting *E. coli* in an ELM under nonoptimal ambient temperature. We demonstrated a genetic circuit for expression of a dark, light-absorptive pigment below 36°C. At 32°C, dense patches of *E. coli* containing this circuit become nearly opaque with black pigment, allowing them to warm above background and grow in both area and thickness at a greater rate than unpigmented patches. Conversely at 42°C, patches remain translucent, avoiding further overheating and growing to a greater thickness than chemically-induced pigmented patches. Because *E.coli-based* ELMs rely on *E. coli* growth and protein production to form the material, as well as confer engineered properties, such as environmental sensing, to the material, long-term exposure to temperatures below or above 37°C will be detrimental to their function. Our temperature switch genetic construct is in principle compatible with any cloning strain of *E. coli* with the *lacZΔM15* genotype, allowing it to be used in existing *E. coli*-based ELMs.

While this study provides a proof of concept for temperature self-regulation, additional work is needed to improve the kinetics of the temperature switch. Because our system does not include a mechanism for degradation of the pigment complex, even brief low-temperature excursions allow pigment to accumulate in the ELM. While protective in the case of long-term exposure to low temperature, this could cause overheating once the ambient temperature reaches 37°C or above. To avoid producing pigment at night in climates or seasons where day-time temperatures exceed 37°C, but night-time temperatures drop below, our circuit could be modified to incorporate a light sensor, such as a phytochrome^[19]^, using an AND gate^[20]^, turning on pigment production only with both light and cold temperature. In addition, to accelerate the transition from dark to translucent in response to a change in environment from low to high temperature, an enzyme cassette for biodegradation of the esculetin-based pigment could be developed from coumarin degradation pathways found in soil bacteria^[21]^. To increase protection from heat, this degradation cassette, or a cassette encoding light-scattering proteins such as gas vesicles ^[22,23]^, could be turned on at high temperature in place of the GFP marker currently incorporated in our construct, clearing pigment from the ELM or scattering incoming light. With our thermal bioswitch circuits and these future improvements, ELMs will gain the ability to harness sunlight to keep warm or stay cool.

## METHODS

### Plasmid construction and molecular biology

All plasmids were designed using SnapGene (GSL Biotech) and assembled via reagents from New England Biolabs for KLD mutagenesis (E0554S) or HiFi Assembly (E2621L). After assembly, constructs were transformed into NEB Turbo (C2984I) *E. coli* for growth and plasmid preparation. Integrated DNA Technologies synthesized all PCR primers. TlpA36 and mWasabi^[24]^ (GFP) were obtained from our previous work^[18]^. LacZα was tagged at the C-terminus with the AAV ssrA tag^[25]^ (amino acid sequence AANDENYAAAV). The weak terminator in the temperature switch construct is Part:BBa_B1002^[26]^. The weak RBS in the temperature switch construct is RBSF^[27]^. Plasmids were transformed into DH10B *E. coli* (ThermoFisher) for downstream experiments. Gene circuit diagrams were created using the DNAplotlib^[28]^ library in Python.

### Visible light absorbance assays

1 mL cultures of LB medium (Sigma) with 100 μg/mL ampicillin were inoculated with a single colony per culture and grown at 38°C, 250 rpm for 18.5 h. 30 μL were diluted into 2 mL LB with 300 μg/mL 3,4-cyclohexenoesculetin-β-D-galactopyranoside (S-gal) (Sigma), 500 μg/mL ferric ammonium citrate (Sigma), 100 μM IPTG, 100 μg/mL ampicillin and propagated at 38°C, 250 rpm for 90 minutes. The cultures were dispensed in 100 μL aliquots into 8-well PCR strips (Bio-Rad) and incubated for 24 h in a thermal gradient using a DNA Engine Tetrad 2 Peltier Thermal Cycler (Bio-Rad) with the lid set to 50°C. PCR strips were imaged using a ChemiDoc MP Gel Imaging System (Bio-Rad). Contrast was adjusted using ImageJ software^[29]^. 90 μL were transferred into 96-well plates (Corning 3631) and absorbance spectra were measured using a Spark multi-mode microplate reader (Tecan). Data analysis was performed using custom Python scripts.

### Preparation of dense patches of E. coli

2 mL cultures of LB medium (Sigma) with 100 μg/mL ampicillin (and 25 μM IPTG for pigmented controls) were inoculated with a single colony per culture and grown at 38°C, 250 rpm for 18.5 h. Nalgene Rapid-Flow sterile disposable filter units (cellulose nitrate, 75 mm diameter, 0.2 μm pore size) (ThermoFisher 450-0020) were rinsed with 1x PBS (Corning) and track-etched polycarbonate membranes (25 mm diameter, 0.2 μm pore size) (Sartorius 23007 or Whatman Nuclepore 10417006) were placed on the filter surface. 200 μL bacterial culture was dispensed onto each membrane and vacuum was applied to the filter unit to coat the cells onto the membrane.

### Growth of E. coli patches in ELM-like conditions

90 mL (32°C experiments) or 100 mL (42°C experiments) of LB medium (Sigma) with 75 μg/mL S-gal, 125 μg/mL ferric ammonium citrate, 25 μM IPTG, 100 μg/mL ampicillin (and 0.1% D-glucose for 42°C experiments) was added to approximately 0.7 g melamine foam (Amazon) in a petri dish until foam was saturated. For 42°C experiments, saturated foam was prewarmed using a heat lamp. Patches of *E. coli* coated on polycarbonate membranes were placed onto the surface of the saturated foam. Patches were incubated either in a custom illuminated incubator or in a conventional incubator for 72 h (32°C experiments) or 48 h (42°C experiments). At 24 h intervals, petri dishes were weighed to determine the amount of evaporation and the equivalent volume of LB with 75 μg/mL S-gal, 125 μg/mL ferric ammonium citrate, 25 μM IPTG, 100 μg/mL ampicillin (and 0.1% D-glucose for 42°C experiments) at room temperature (for 32°C experiments) or prewarmed to 42°C (for 42°C experiments) was added.

### Design and construction of illuminated incubator with in situ temperature monitoring

The enclosure of the illuminated incubator is an 203.2 mm x 254 mm x 180.34 mm aluminum box with a lid consisting of a 3.175 mm layer of glass (IR-filtering) and a 1.5 mm layer of acrylic (UV-filtering) held in a 3D-printed polylactic acid (PLA) collar. Weatherproofing foam was used to provide a tight fit between the lid and the box. The sample holder is located 120 mm from the 100 W LED grid light source (Mifxion), receiving an average light flux of 120000 lx. The light source is plugged into a power relay (Digital Loggers). The light source heats the interior of the chamber; the temperature is adjusted by adding or removing insulation.

A MLX90640 32 x 24 thermal IR sensor array (Grove or Adafruit) is attached to a SG92R micro servo motor (Tower Pro) via 3D-printed PLA fittings reinforced with wooden craft sticks for monitoring the samples. MCP9808 temperature sensors (Adafruit) were used to monitor the air temperature inside and outside the sample enclosure. All sensors, the servo motor, and the power relay were connected to a Feather M4 Express microcontroller (Adafruit) for data acquisition and control. A custom Bokeh application was used to interface with the microcontroller.

### Imaging and analysis of E. coli patches

*E. coli* patches on polycarbonate membranes were transferred from melamine foam growth substrate to 20 mL PBS agar plates for imaging.

White light transillumination imaging was performed using a ChemiDoc MP Gel Imaging System (Bio-Rad). *E. coli* patch area and pixel intensity (normalized to polycarbonate membrane background and opaque black plastic) was measured by using the Canny edge detector algorithm for image segmentation via custom Python scripts.

Optical coherence tomography was performed using a Ganymede 210 Series Spectral Domain OCT Imaging System with an OCTP-900 scanner and 8 μm lateral resolution scan lens (Thorlabs). Cross-sectional images with a 10 mm / 2560 pixel (x) by 1 mm / 491 pixel (z) field of view were acquired at 5.5 kHz A-scan rate with 7 B-scan averages and 5 A-scan averages per acquisition. The despeckle filter in the ThorImageOCT software (Thorlabs) was applied before exporting. *E. coli* patch thickness was measured using a custom Python script. Briefly, after applying a Gaussian filter, the 3 narrowest peaks with height and width fulfilling adjustable thresholds are detected in each column and arranged by distance from the top of the image to correspond with the top of the patch, the top of the polycarbonate membrane, and the bottom of the polycarbonate membrane. The vectors for each interface are Hampel filtered to reduce outliers due to imaging artifacts. The thickness is defined as the distance from the top of the patch to the top of the polycarbonate membrane. The mean thickness and standard deviation of each patch between x = 3.5 mm and x = 9.0 mm (avoiding the edges of the patch) was calculated. Then, the mean for each construct, illumination condition, and temperature was calculated, propagating the standard deviation to calculate the standard error of the mean.

## AUTHOR CONTRIBUTIONS

L.L.X. and M.G.S. conceived the study. L.L.X. and M.A.G. planned and performed experiments. L. L.X. analyzed data. J. A. K. provided input on research design and data interpretation. L.L.X. and M. G.S. wrote the manuscript with input from all other authors. M.G.S. and J.A.K. supervised the research.

## ACKNOWLEDGEMENTS

The authors thank Hanwei Liu, David Tirrell, Priya Chittur, and Seunghyun Sim for helpful discussions about engineered living materials, as well as Red Lhota, Robert Learsch, and Justin Bois for assistance with instrument design. This research was supported by the Defense Advanced Research Project Agency (HR0011-17-2-0037 to M.G.S. and J.A.K.), the Institute for Collaborative Biotechnologies (W911NF-19-D-0001 to M.G.S.), the Jacobs Institute for Molecular Engineering for Medicine (to J.A.K.), and the Elizabeth W. Gilloon Chair (to J.A.K.). L.L.X. was supported by the NSF Graduate Research Fellowship Program. M.A.G. was supported by the NIH MBRS Research Initiative for Scientific Enhancement Program. M.G.S. is an Investigator of the Howard Hughes Medical Institute. Related research in the Shapiro lab is supported by the David and Lucile Packard Foundation and the Dreyfus Foundation.

## REFERENCES

[1] P. Q. Nguyen, N.-M. D. Courchesne, A. Duraj-Thatte, P. Praveschotinunt, N. S. Joshi, Adv. Mater. 2018, 30, 1704847.

[2] C. Gilbert, T. Ellis, ACS Synth. Biol. 2019, 8, 1.

[3] S. Liu, Xu, Front. Sens. 2020, 1, DOI 10.3389/fsens.2020.586300.

[4] L. K. Rivera-Tarazona, T. Shukla, K. A. Singh, A. K. Gaharwar, Z. T. Campbell, T. H. Ware, Adv. Funct. Mater. 2022, 32, 2106843.

[5] C. M. Heveran, S. L. Williams, J. Qiu, J. Artier, M. H. Hubler, S. M. Cook, J. C. Cameron, W. V. Srubar, Matter 2020, 2, 481.

[6] J. Huang, S. Liu, C. Zhang, X. Wang, J. Pu, F. Ba, S. Xue, H. Ye, T. Zhao, K. Li, Y. Wang, J. Zhang, L. Wang, C. Fan, T. K. Lu, C. Zhong, Nat. Chem. Biol. 2019, 15, 34.

[7] L. M. González, N. Mukhitov, C. A. Voigt, Nat. Chem. Biol. 2020, 16, 126.

[8] K. C. Heyde, F. Y. Scott, S.-H. Paek, R. Zhang, W. C. Ruder, JoVE J. Vis. Exp. 2017, e55300.

[9] A. M. Duraj-Thatte, N.-M. D. Courchesne, P. Praveschotinunt, J. Rutledge, Y. Lee, J. M. Karp, N. S. Joshi, Adv. Mater. 2019, 31, 1901826.

[10] S. Guo, E. Dubuc, Y. Rave, M. Verhagen, S. A. E. Twisk, T. van der Hek, G. J. M. Oerlemans, M. C. M. van den Oetelaar, L. S. van Hazendonk, M. Brüls, B. V. Eijkens, P. L. Joostens, S. R. Keij, W. Xing, M. Nijs, J. Stalpers, M. Sharma, M. Gerth, R. J. E. A. Boonen, K. Verduin, M. Merkx, I. K. Voets, T. F. A. de Greef, ACS Synth. Biol. 2020, 9, 475.

[11] Z. D. Blount, eLife n.d., 4, e05826.

[12] M. D. Winfield, E. A. Groisman, Appl. Environ. Microbiol. 2003, 69, 3687.

[13] J. D. van Elsas, A. V. Semenov, R. Costa, J. T. Trevors, ISME J. 2011, 5, 173.

[14] M. K. Shaw, A. G. Marr, J. L. Ingraham, J. Bacteriol. 1971, 105, 683.

[15] R. J. Broeze, C. J. Solomon, D. H. Pope, J. Bacteriol. 1978, 134, 861.

[16] B. Rudolph, K. M. Gebendorfer, J. Buchner, J. Winter, J. Biol. Chem. 2010, 285, 19029.

[17] C. A. Voigt, Synthetic Biology: Methods for Part/Device Characterization and Chassis Engineering, Academic Press, 2011.

[18] D. I. Piraner, M. H. Abedi, B. A. Moser, A. Lee-Gosselin, M. G. Shapiro, Nat. Chem. Biol. 2017, 13, 75.

[19] A. Levskaya, A. A. Chevalier, J. J. Tabor, Z. B. Simpson, L. A. Lavery, M. Levy, E. A. Davidson, A. Scouras, A. D. Ellington, E. M. Marcotte, C. A. Voigt, Nature 2005, 438, 441.

[20] J. C. Anderson, C. A. Voigt, A. P. Arkin, Mol. Syst. Biol. 2007, 3, 133.

[21] A. Krikštaponis, R. Meškys, Mol. J. Synth. Chem. Nat. Prod. Chem. 2018, 23, 2613.

[22] A. E. Walsby, Microbiol. Rev. 1994, 58, 94.

[23] R. W. Bourdeau, A. Lee-Gosselin, A. Lakshmanan, A. Farhadi, S. R. Kumar, S. P. Nety, M. G. Shapiro, Nature 2018, 553, 86.

[24] H. Ai, S. G. Olenych, P. Wong, M. W. Davidson, R. E. Campbell, BMC Biol. 2008, 6, 13.

[25] “Part:BBa_I11012,” can be found under http://parts.igem.org/Part:BBa_I11012, n.d.

[26] “Part:BBa_B1002,” can be found under http://parts.igem.org/Part:BBa_B1002, n.d.

[27] A. Levin-Karp, U. Barenholz, T. Bareia, M. Dayagi, L. Zelcbuch, N. Antonovsky, E. Noor, R. Milo, ACS Synth. Biol. 2013, 2, 327.

[28] B. S. Der, E. Glassey, B. A. Bartley, C. Enghuus, D. B. Goodman, D. B. Gordon, C. A. Voigt, T. E. Gorochowski, ACS Synth. Biol. 2017, 6, 1115.

[29] C. A. Schneider, W. S. Rasband, K. W. Eliceiri, Nat. Methods 2012, 9, 671.

