## Supplementary Material for "Self-regulating living material with temperature-dependent light absorption"

***Engineering thermally self-regulating bacteria with temperature-dependent light absorption***

[1] Division of Engineering and Applied Sciences
[2] Division of Chemistry and Chemical Engineering
[3] Howard Hughes Medical Institute
California Institute of Technology
Pasadena, CA USA 91125

**Table of contents**

Supplementary Tables S1-S2
Supplementary Figures S1-S6

### SUPPLEMENTARY TABLES S1-S2

**Supplementary Table S1.** Genetic constructs used in this study.

| Plasmid | Purpose | Transcriptional Regulator(s) | Output Gene Product(s) |
|---|---|---|---|
| pTSwitch-LacZα | temperature switch | TlpA36, CI | LacZα, mWasabi |
| pTlpA36-wasabi | unpigmented control | TlpA36 | mWasabi |
| pTrcLacZα | pigmented control | LacIq | LacZα |

**Supplementary Table S2.** Sequences of DNA parts used in this study.

| Type | Name | Sequence |
|---|---|---|
| promoter | PTlpA | TTTAATTTGTTTGTTAGTTAGTTTATTTGTTGGTTTGTTTGTGTTATA ATAT |
| promoter | PR | GTGCGTGTTGACTATTTTACCTCTGGCGGTGATAATGGTTGCATGTACTAAG GAGGTTG |
| promoter | PL | AACCATCTGCGGTGATAAATTATCTCTGGCGGTGTTGACATAAATACCACTG GCGGTGATACTGAGCACATCAGCAGG |
| promoter | PTrc | TTGACAATTAATCATCCGGCTCGTATAATGTGTGGAATTGTGAGCGGATAAC AATT |
| degradation tag | AAV ssrA tag | GCTGCTAACGACGAAAACTACGCTGACGCTTCT |
| terminator | T7 | CTAGCATAACCCCTTGGGGCCTCTAAACGGGTCTTGAGGGGTTTTTTG |
| terminator | Part:BBa_B1002 | CGCAAAAAACCCCGCTTCGGCGGGGTTTTTTCGC |
| RBS | RBSF | CACCATACACTG |
| gene | LacZa | CATGATTACGGATTCACTGGCCGTCGTTTTACAACGTCGTGACTGGGAAAAC CCTGGCGTTACCCAACTTAATCGCCTTGCAGCACATCCCCCTTTCGCCAGCT GGCGTAATAGCGAAGAGGCCCGCACCGATCGCCCTTCCCAACAGTTGCGCA GCCTGAATGGCGAATGGCGCTTTGCCTGGTTTCCGGCACCAGAAGCGGTGC CGGAAAGCTGGCTGGAG |
| gene | mWasabi | ATGGTGAGCAAGGGCGAGGAGACCACAATGGGCGTAATCAAGCCCGACATG AAGATCAAGCTGAAGATGGAGGGCAACGTGAATGGCCACGCCTTCGTGATCG AGGGCGAGGGCGAGGGCAAGCCCTACGACGGCACCAACACCATCAACCTGG AGGTGAAGGAGGGAGCCCCCCTGCCCTTCTCCTACGACATTCTGACCACCGC GTTCAGTTACGGCAACAGGGCCTTCACCAAGTACCCCGACGACATCCCCAAC TACTTCAAGCAGTCCTTCCCCGAGGGCTACTCTTGGGAGCGCACCATGACCT TCGAGGACAAGGGCATCGTGAAGGTGAAGTCCGACATCTCCATGGAGGAGG ACTCCTTCATCTACGAGATACACCTCAAGGGCGAGAACTTCCCCCCCAACGG CCCCGTGATGCAGAAGGAGACCACCGGCTGGGACGCCTCCACCGAGAGGAT GTACGTGCGCGACGGCGTGCTGAAGGGCGACGTCAAGATGAAGCTGCTGCT GGAGGGCGGCGGCCACCACCGCGTTGACTTCAAGACCATCTACAGGGCCAA GAAGGCGGTGAAGCTGCCCGACTATCACTTTGTGGACCACCGCATCGAGATC CTGAACCACGACAAGGACTACAACAAGGTGACCGTTTACGAGATCGCCGTGG CCCGCAACTCCACCGACGGCATGGACGAGCTGTACAAGGGC |
| gene | TlpA36 | ATGCGTCCGGCGACATACGAACCAGAACAGATTATTGAAGCAGGGCTGGCCC TGCAGGCTGAAGGACGGAATATCACCGGGTTCGCACTACGTAACCAGGTGG GTGGCGGCAATCCGACACGTCTCCGCCAGATATGGGACGAATACCAGGCTT CACAGAGCACGGTCGTCACTGAACTCGTTGCCGAGCTGCCAGTGGAAGTGG CTGAAGAAGTGAAGGCCGTCTCCGCCGCGCTGTCCGAACGCATCACCCAGC TGGCGACAGAACTGAATGACAAGGCGGTCCGGGCTGCAGAACGCCGGGTTG CGGAAGTCACGCGTGCTGCCGGTGAACAGACCGCACAGGCAGAGCGGGAGC TGGCCGACGCCGCGCAGACAGTCGACGACCTGGAAGAAAAACTGGTTGAACT GCAGGACAGATATGACAGTTTGACGCTGGCGCTGGAGTCAGAACGTTCACT GCGTCAGCAGCATGATGTGGAGATGGCCCAGCTGAAAGAGCGTCTTGCGGC CGCTGAAGAGAATACCCGTCAGCGAGAGGAACGGTATCAGGAGCAGAGGAC AGTGCTGCAGGATGCGCTTAATGCGGAGCAGGCACAGCACATAAACACGCG GGAAGACCAGCAGAAACGACTGGAGCAAATTTCTGCCGAAGCTAATGCGCGT ACAGAAGAACTGAAGTCTGAACGCGATAAAGTCAATACTCTCCTTACCCGCC TTGAATCGCAGGAAAATGCGCTGGCCTCAGAACGTCAGCAGCATCTGGCCAC CCGCGAAACGCTGCAGCAACGCCTCGAGCAGGCCATCGCTGACACGCAGGC GCGCGCCGGTGAGATTGCACTTGAACGTGACAGAGTCAGCAGCCTCACCGC AAGGCTGGAATCGCAGGAAAAGGCCTCCTCGGAGCAACTGGTGCGTATGGG CAGTGAAATAGCCAGTCTGACAGAGCGTTGCACACAGCTGGAAAACCAGCGT GATGATGCCCGTCTGGAGACGATGGGGGAGAAAGAAACGGTCGCGGCACTG CGTGGTGAGGCTGAAGCCCTGAAGCGTCAGAACCAGTCACTGATGGCGGCG CTTTCAGGCAATAAACAGACCGGTGGCCAGAATGCGT |

| | | |
|---|---|---|
| gene | CI | ATGAGCACAAAAAAGAAACCATTAACACAAGAGCAGCTTGAGGACGCACGTC<br>GCCTTAAAGCAATTTATGAAAAAAAGAAAAATGAACTTGGCTTATCCCAGGAA<br>TCTGTCGCAGACAAGATGGGGATGGGGCAGTCAGGCGTTGGTGCTTTATTT<br>AATGGCATCAATGCATTAAATGCTTATAACGCCGCATTGCTTGCAAAAATTCT<br>CAAAGTTAGCGTTGAAGAATTTAGCCCTTCAATCGCCAGAGAAATCTACGAG<br>ATGTATGAAGCGGTTAGTATGCAGCCGTCACTTAGAAGTGAGTATGAGTACC<br>CTGTTTTTTCTCATGTTCAGGCAGGGATGTTCTCACCTGAGCTTAGAACCTT<br>TACCAAAGGTGATGCGGAGAGATGGGTAAGCACAACCAAAAAAGCCAGTGAT<br>TCTGCATTCTGGCTTGAGGTTGAAGGTAATTCCATGACCGCACCAACAGGCT<br>CCAAGCCAAGCTTTCCTGACGGAATGTTAATTCTCGTTGACCCTGAGCAGGC<br>TGTTGAGCCAGGTGATTTCTGCATAGCCAGACTTGGGGGTGATGAGTTTAC<br>CTTCAAGAAACTGATCAGGGATAGCGGTCAGGTGTTTTTACAACCACTAAAC<br>CCACAGTACCCAATGATCCCATGCAATGAGAGTTGTTCCGTTGTGGGGAAAG<br>TTATCGCTAGTCAGTGGCCTGAAGAGACGTTTGGCTGA |

### SUPPLEMENTARY FIGURES S1-S6

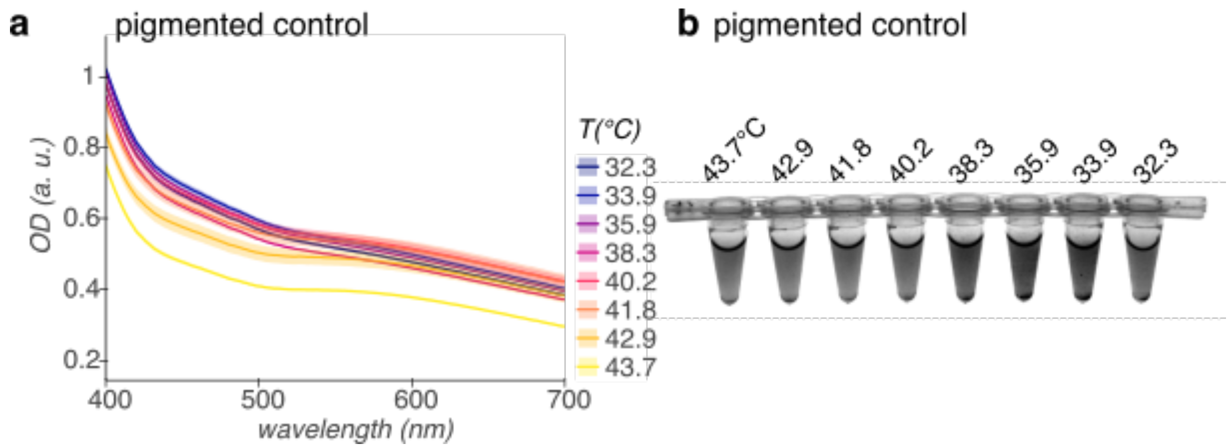


**Supplementary figure S1.** Visible light absorption spectra (**a**) and representative white light transillumination image (**b**) of cultures of *E. coli* containing a pigmented control construct encoding IPTG-inducible LacZα after 24 h growth in pigment-induction media at temperatures ranging from 43.7°C to 32.3°C. $n$ = 2 biological replicates; shading represents +/- standard error of the mean.

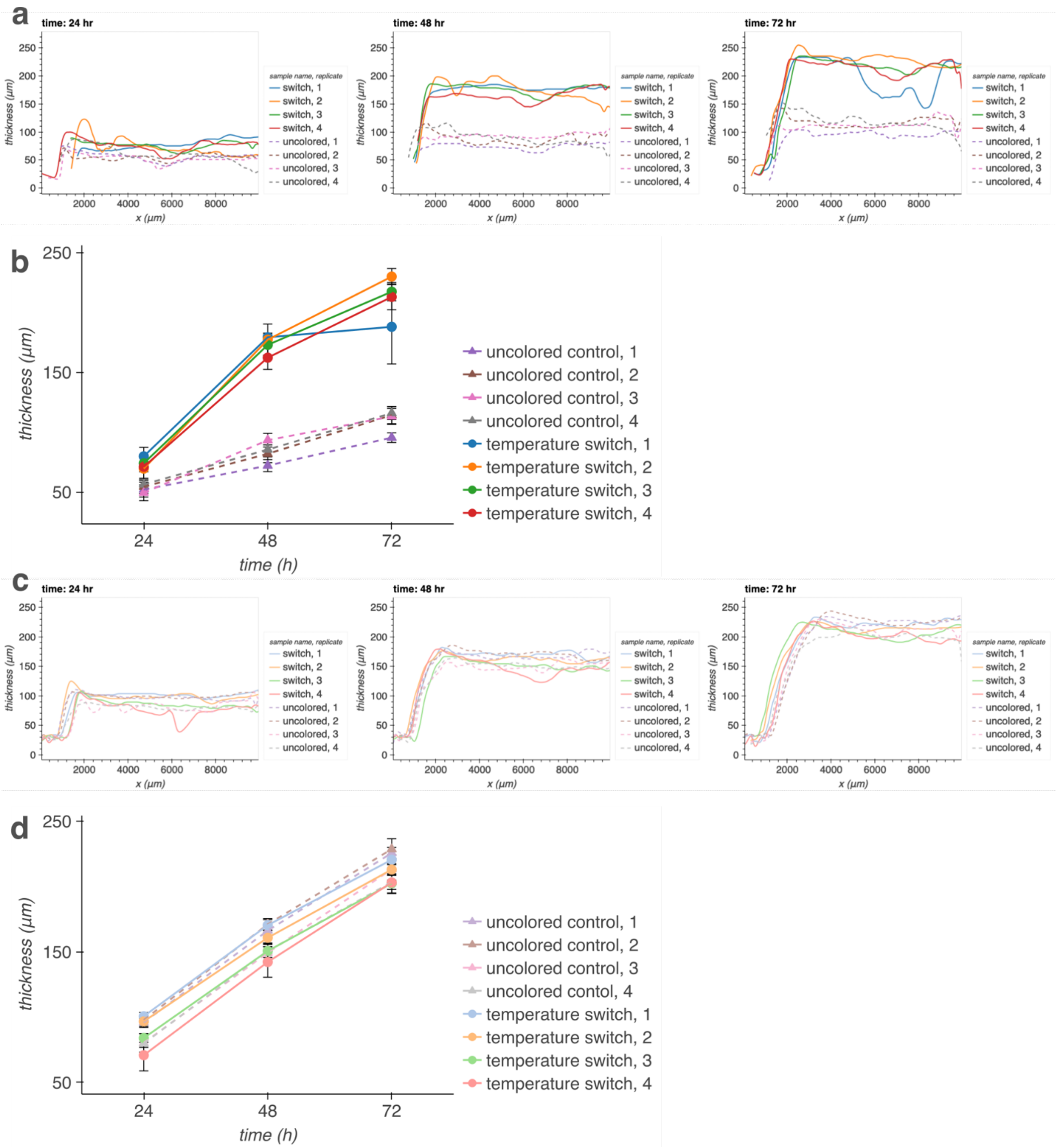


**Supplementary figure S2.** (**a, c**) Thicknesses over time measured across full OCT cross-section for each patch of *E. coli* containing either the temperature switch construct or an unpigmented control construct encoding heat-inducible GFP, grown at 32°C with (**a**) or without (**c**) illumination. (**b, d**) Mean thickness and standard deviation between x = 3.5 mm and x = 9.0 mm (avoiding the edges of the patch) of each patch, grown with (**b**) or without (**d**) illumination over time.

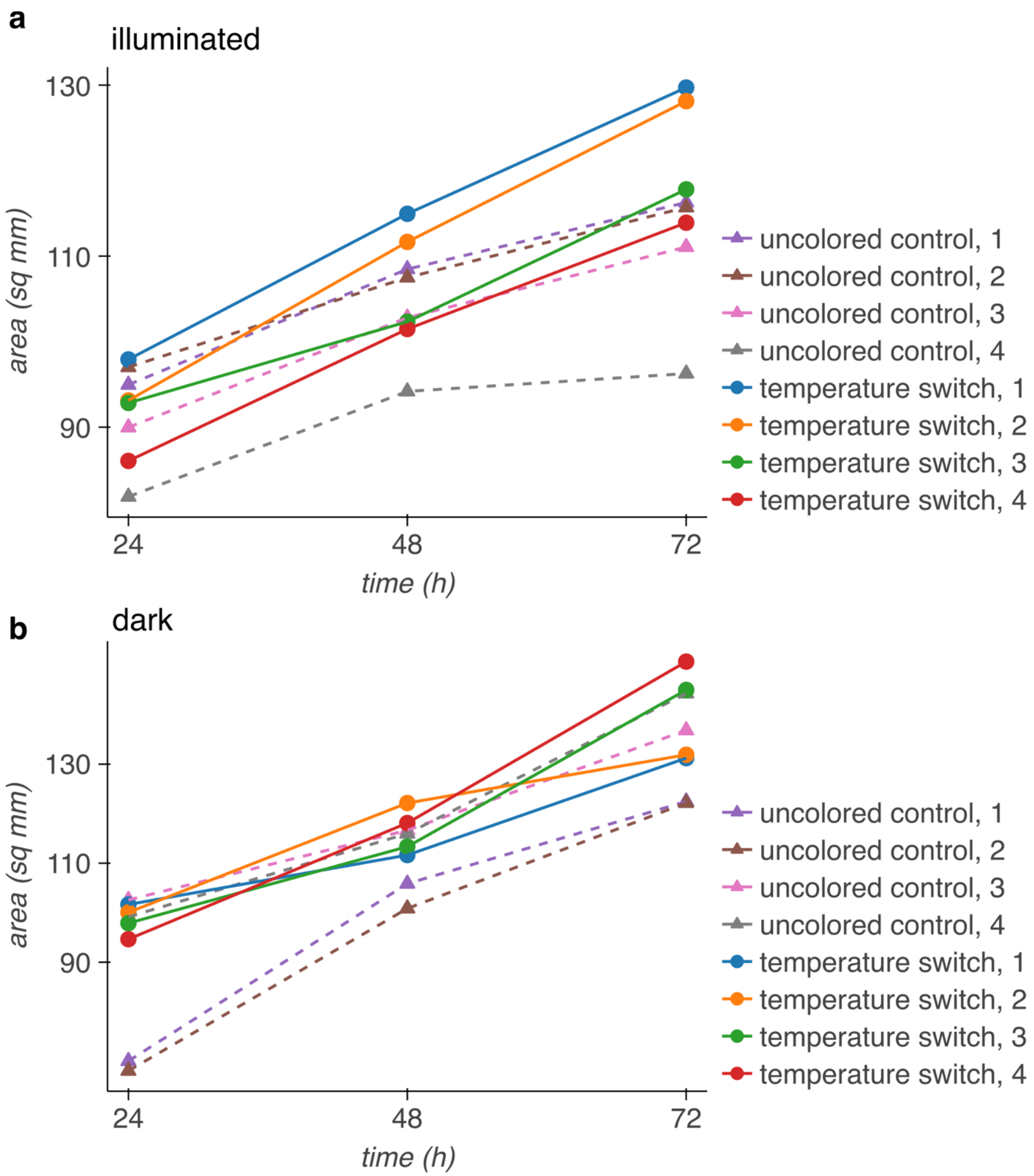


**Supplementary figure S3.** Area of patches of *E. coli* containing either the temperature switch construct or an unpigmented control construct encoding heat-inducible GFP, grown at 32°C with (**a**) or without (**b**) illumination over time.

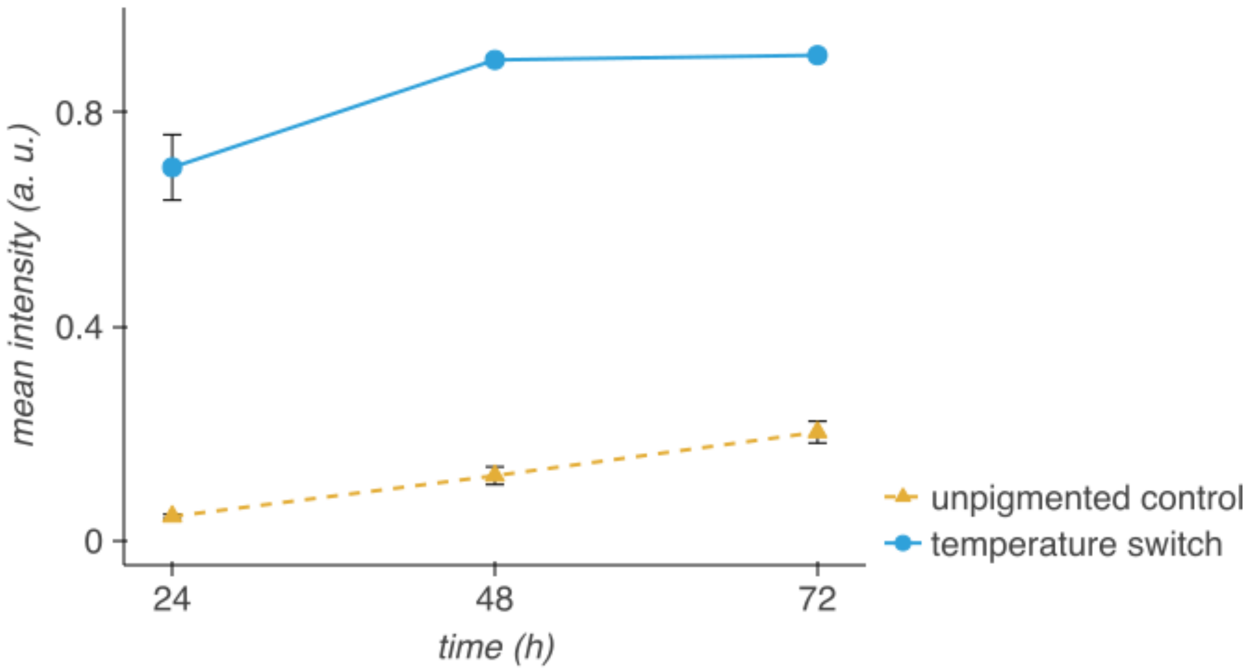


**Supplementary figure S4.** Mean pixel intensity of patches of *E. coli* containing either the temperature switch construct or a n unpigmented control construct encoding heat-inducible GFP, grown at 32°C under illumination, over time. Image was normalized so that the polycarbonate membranes have a mean intensity of 0 and opaque black plastic has a mean intensity of 1.

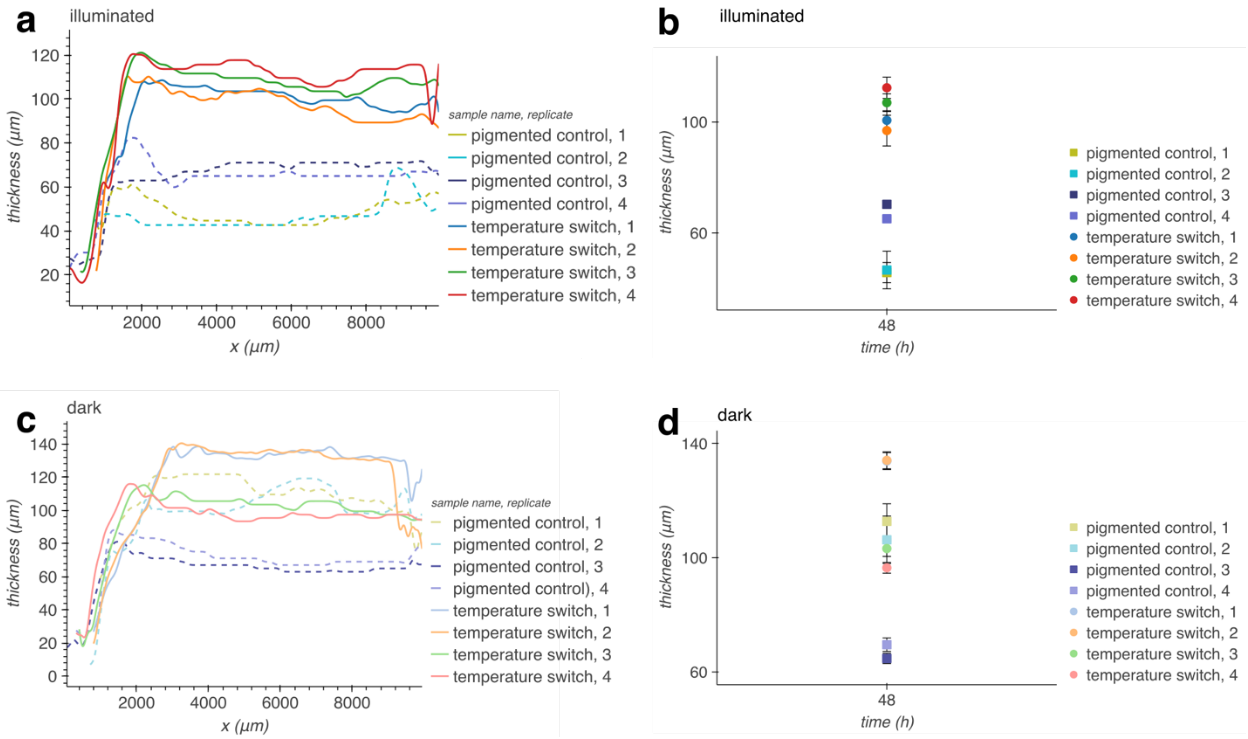


**Supplementary figure S5.** (**a, c**) Thicknesses over time measured across full OCT cross-section for each patch of *E. coli* containing either the temperature switch construct or a pigmented control construct encoding IPTG-inducible LacZα, grown at 42°C with (**a**) or without (**c**) illumination. (**b, d**) Mean thickness and standard deviation between x = 3.5 mm and x = 9.0 mm (avoiding the edges of the patch) of each patch, grown with (**b**) or without (**d**) illumination over time.

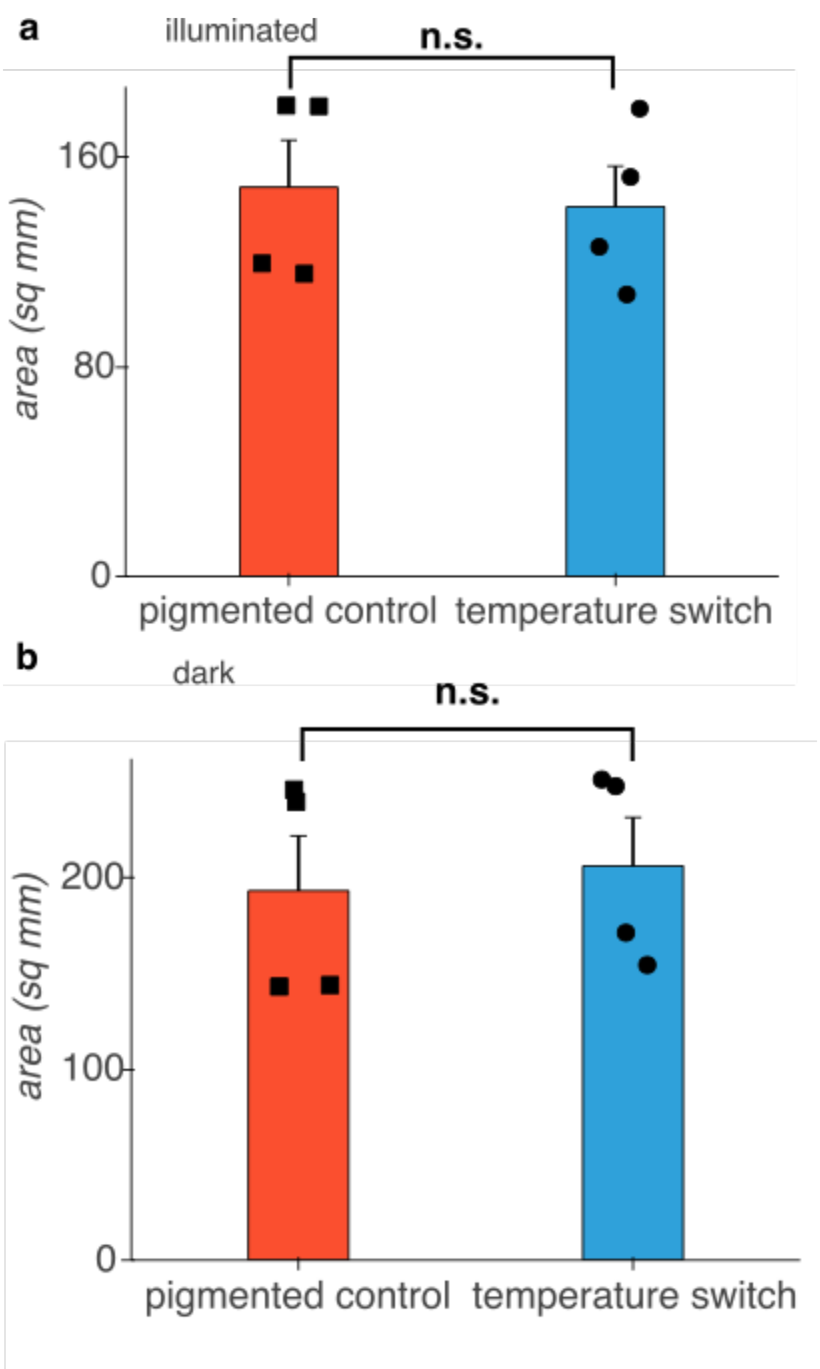


**Supplementary figure S6.** Area of patches of *E. coli* containing either the temperature switch construct or a pigmented control construct encoding IPTG-inducible LacZα, grown at 42°C with (**a**) or without (**b**) illumination for 48 h.
